# Reference data based insights expand understanding of human metabolomes

**DOI:** 10.1101/2020.07.08.194159

**Authors:** Julia M. Gauglitz, Wout Bittremieux, Candace L. Williams, Kelly C. Weldon, Morgan Panitchpakdi, Francesca Di Ottavio, Christine M. Aceves, Elizabeth Brown, Nicole C. Sikora, Alan K. Jarmusch, Cameron Martino, Anupriya Tripathi, Erfan Sayyari, Justin P. Shaffer, Roxana Coras, Fernando Vargas, Lindsay DeRight Goldasich, Tara Schwartz, MacKenzie Bryant, Gregory Humphrey, Abigail J. Johnson, Katharina Spengler, Pedro Belda-Ferre, Edgar Diaz, Daniel McDonald, Qiyun Zhu, Dominic S. Nguyen, Emmanuel O. Elijah, Mingxun Wang, Clarisse Marotz, Kate E. Sprecher, Daniela Vargas Robles, Dana Withrow, Gail Ackermann, Lourdes Herrera, Barry J. Bradford, Lucas Maciel Mauriz Marques, Juliano Geraldo Amaral, Rodrigo Moreira Silva, Flávio Protaso Veras, Thiago Mattar Cunha, Rene Donizeti Ribeiro Oliveira, Paulo Louzada-Junior, Robert H. Mills, Douglas Galasko, Parambir S. Dulai, Curt Wittenberg, David J. Gonzalez, Robert Terkeltaub, Megan M. Doty, Jae H. Kim, Kyung E. Rhee, Julia Beauchamp-Walters, Kenneth P. Wright, Maria Gloria Dominguez-Bello, Mark Manary, Michelli F. Oliveira, Brigid S. Boland, Norberto Peporine Lopes, Monica Guma, Austin D. Swafford, Rachel J. Dutton, Rob Knight, Pieter C. Dorrestein

## Abstract

The human metabolome has remained largely unknown, with most studies annotating ∼10% of features. In nucleic acid sequencing, annotating transcripts by source has proven essential for understanding gene function. Here we generalize this concept to stool, plasma, urine and other human metabolomes, discovering that food-based annotations increase the interpreted fraction of molecular features 7-fold, providing a general framework for expanding the interpretability of human metabolomic “dark matter.”

## Introduction

In 2016, typical MS/MS-based untargeted metabolomics studies annotated only ∼2% of molecules based on matches against spectral libraries, leaving the rest of the sample as metabolomic “dark matter.” The capture of community knowledge, accumulating public reference MS/MS spectra over the past four years, has increased this baseline ∼2.5-fold within the global natural product social molecular networking (GNPS) infrastructure (Wang et al., 2016). This growth has been even more dramatic for data from commonly-studied specimen types such as human stool and plasma: 10.1 +/- 4.4% of MS/MS features now match to a reference MS/MS spectrum [1% FDR (Scheubert et al., 2017), n = 30, average number of unique MS/MS spectra is 12,889/dataset]. However, despite these advances, the vast majority of detectable spectra lack any annotation.

This situation for MS/MS spectra is in sharp contrast to the interpretability of uncharacterized portions of the human genome. For example, reference data sets for gene expression, such as expressed sequence tags (an early form of RNASeq), enable the sequencing of “dark matter,” as opposed to monitoring the expression of a single curated gene. Such methods have significantly improved interpretation by annotating genes not directly by function, but rather by source (developmental stage, tissue location, organism-level, phenotype, etc.) (Bono, 2020; Ono et al., 2017). Interpretation based on source has been very important for metagenomics and metatranscriptomics, increasing our understanding of the structure and function of complex communities by leveraging matches between genes or transcripts of known and unknown origin via publicly available databases.

Annotation of chemicals, based on their source within publicly available complex reference samples that use controlled metadata vocabularies, has not been applied to metabolomics for several reasons. First, standards for annotation of molecules that are used to create spectral libraries have been based on availability of individual pure, typically commercially available, standards, and structural considerations such as presence of specific moieties. Many molecules are observed as multiple different ion forms, such as adducts, in-source fragments, and multimers. Current spectral libraries do not contain all possible ion forms of those molecules, and typically only the protonated form (Schmid et al., 2020; Vinaixa et al., 2016), because reference standards that run in a highly purified state that biases towards detection extraction of only specific data on specific ion forms. These forms are often different from the ions associated with the same molecule present in an extract from a biological matrix (e.g. proton vs sodium or even multiple sodium and potassium adducts), which then cannot be matched because the relevant spectra are not in the database. Second, on average, 5–10% of untargeted metabolomics data can be annotated from spectral libraries: the remaining 90+% are unassignable “dark matter” in metabolomics, especially when obtained from complex matrices such as human samples. Third, large databases of untargeted metabolomics data with consistently annotated provenance with controlled vocabularies have been neither available nor possible to effectively reuse. We recently addressed this latter problem via GNPS (Wang et al., 2016), ReDU (Jarmusch et al., 2019), importing data from MetaboLights into GNPS (Haug et al., 2020), with ReDU-compatible metadata conversion. Finally, the availability of robust scalable analysis infrastructures and algorithms, such as molecular networking, that enable the functional equivalent of reporting of expressed sequence tag/RNASeq analysis, have only recently been introduced for mass spectrometry (Wang et al., 2016; Watrous et al., 2012).

To improve interpretation of otherwise unannotated data from untargeted mass spectrometry experiments, we leverage entire reference data sets with curated ontologies to complement existing spectral libraries of individual molecules. Due to lack of a better term we refer to this approach as interpretive metabolomics in this manuscript, and demonstrate its potential by leveraging the Global FoodOmics MS/MS spectral database, which we have made publicly available on MassIVE. This food reference data set will be key for enabling future insights into human health given the importance of diet and the urgent need to develop additional methods for empirical nutrient and diet assessments to understand acute and chronic human disease (Barabási et al., 2020). We demonstrate that interpretive metabolomics can address these types of knowledge gaps by showing that it not only massively expands the fraction of the data that can be interpreted, but that these new insights can lead to an improved understanding of the diets consumed upon co-analysis of human and food/beverage mass spectral data.

## Results/Discussion

We conjectured that a major source of chemicals detected by metabolomics in human samples originates from foods and beverages. We created “Global FoodOmics” (http://www.globalfoodomics.org) in 2017, which now contains 3,579 food and beverage samples contributed by the community, as outlined in the methods, following in the footsteps of the American Gut and the Earth Microbiome Projects (McDonald et al., 2018; Thompson et al., 2017). The majority of samples were photographed, and a subset were subjected to 16S rRNA profiling (1,511 samples) to characterize the microbial composition, as well as providing information about mitochondria and chloroplast sequences matched by the same primers. Foods were manually classified according to the Earth Microbiome Project Ontology, the USDA Food Composition Database and a modification of the Food and Nutrient Database for Dietary Studies (Johnson et al., 2019; Thompson et al., 2017) (https://ndb.nal.usda.gov/) to allow cross-study compatibility. In total, we report 157 metadata categories that further include a six-level food ontology, as well as fermentation or organic status, land or aquatic origin, country of origin, etc. (**Table S1**). Foods and beverages in Global FoodOmics consist of a range of items, from simple ingredients to prepared meals, as well as animal feed.

A key benefit of interpretive metabolomics is that we consider all different ion forms encountered while collecting the Global FoodOmics dataset. The millions of MS/MS spectra in Global FoodOmics inherently include MS/MS spectra of different ion forms of both known and unknown molecules, and can, therefore, be matched in human biospecimens via direct matching of the MS/MS spectra or by more sophisticated approaches. The similar complexity of the reference and experimental data includes many chemicals that may have uncharacterized behavior, such as unexpected adducts or even multimers made up of different molecules. For the MS/MS spectra that do have annotations, it is possible to leverage GNPS tags to test whether the spectral matches make sense in the context of Global FoodOmics.

Within the GNPS environment, the community can also add tags to each reference spectrum in the spectral library using a controlled vocabulary, including multiple per structure. An InChIKey was included for 4586 of 5455 spectral matches against the reference libraries (∼5% annotation rate at 1% FDR), which yielded 1492 unique structures upon consideration of planar structures. There were 415/1492 structures that had lifestyle tags and “food consumption” is the most frequently reported with 357 entries (86%) (**Figure S1a**) (Bouslimani et al., 2016). Brief descriptive tags provide more detail about the annotation itself, and 1131/1492 structures were annotated with such tags. The most common descriptive tags were in order: “natural product” (790/1131), “food” (576/1131), “human”, “plant”, “natural product_plant”, “plant_angiospermae”, and “drug” (**Figure S1b**). Some of these associations with the category “human” may also be of food origin, such as arachidonoyl carnitine, which is currently only tagged as “human,” but may have a variety of animal-product based food sources. Similarly, the tag “drug” includes annotations such as the antimicrobial agent monensin, which is not tagged as a food molecule, but is consumed with animal products from animals raised using monensin as a growth promoter. Thus the Global FoodOmics reference data capture not only inherently food-derived molecules, but also food-sourced exogenous compounds such as preservatives, growth enhancing substances, antimicrobials, pesticides, and packaging materials. However, because the annotation rate remains low, most of the data remains unused despite the informative tags.

In addition to annotating molecules based on matches to library spectra, spectral matches to the food reference data can be obtained and visualized using MS/MS based molecular networking. When applying this method to both foods and biospecimens in an experimental sleep restriction and circadian misalignment study we observed connectivity of nodes within molecular families representing MS/MS spectra (**Figure 1a,b**). Using spectral libraries the tomatidine molecular family was shown to contain both annotated nodes (level 2 or 3, according to the 2007 metabolomics standards initiative (Sumner et al., 2007) e.g, tomatidine, solasodine and sarsasapogenin (**Figure 1b**), as well as unannotated nodes, which are also observed with molecules occurring within Nightshade (Solanaceae) samples from the Global FoodOmics data set (**Figure 1c**). Sarsasapogenin (**Figure 1c, node 1**) is found in food as well as stool data while the +15.996 Da, the addition of the atom “O”, is only observed in stool data. However, numerous other molecular families (such as **Figure 1c, node 10**) contain no annotation, but do have spectral matches between plasma and foods — in this case features also observed in grape and fermented grape samples. In other cases, a plasma metabolite is annotated and connected to unannotated compounds found within the food reference samples (**Figure 1c, nodes 11-14**). These examples highlight how molecular networking can be used to propagate potential metabolism. How potential metabolism can be inferred with molecular networking is explained in (Quinn et al., 2017) and (Aron et al., 2020).

**Figure 1.**
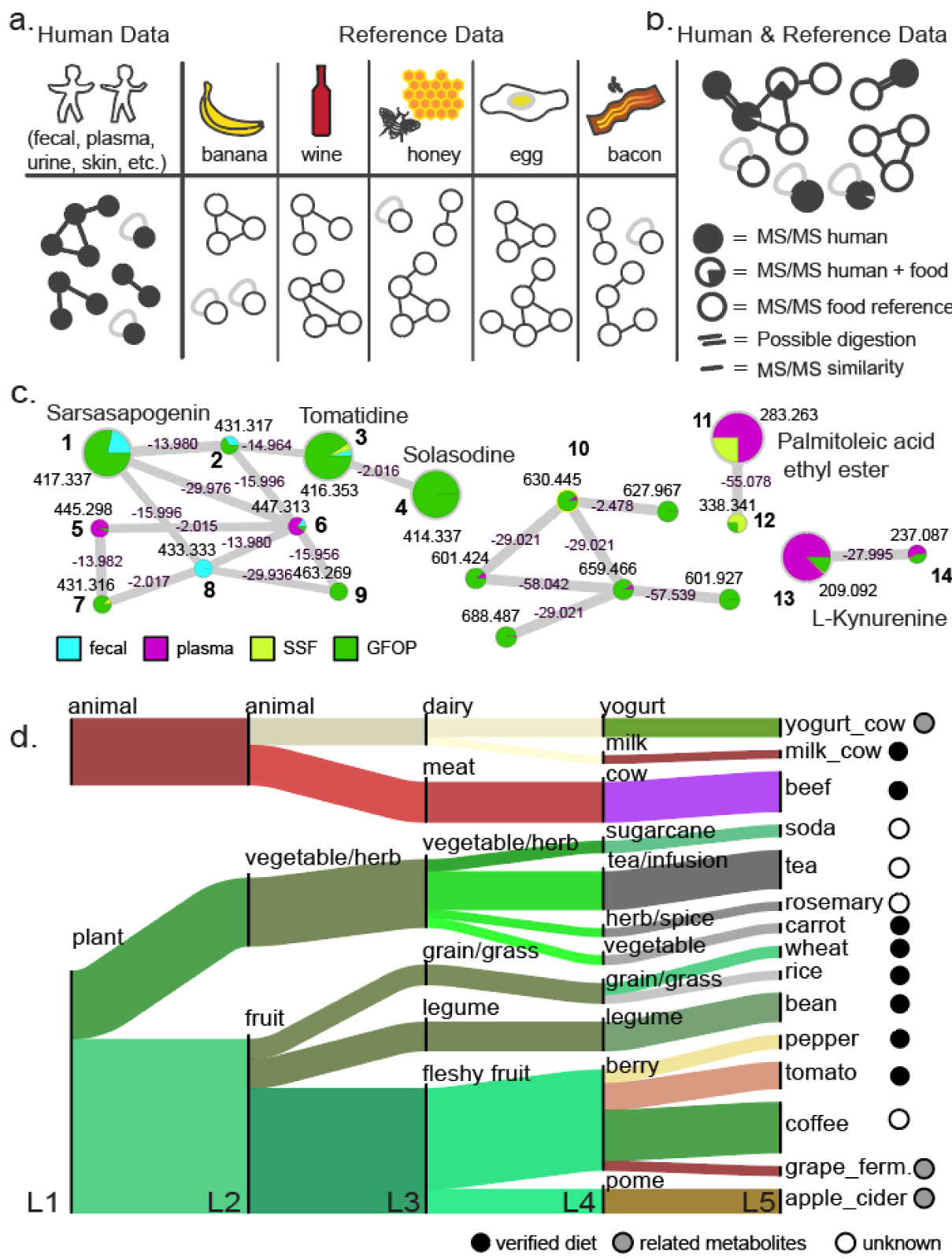
The concept of interpretive metabolomics leveraging reference data sets. **a**. A schematic overview of human data and reference data (e.g. data from food items) as molecular families from independent data sets that are used in b. **b**. A schematic representation when reference data is co-networked with human metabolomics data. Each node represents a unique MS/MS spectrum. **c**. Experimentally observed molecular families (sub-networks) generated from the co-analysis of stool (light blue) and plasma (magenta) data from a sleep restriction and circadian misalignment study with the Global FoodOmics reference dataset (green). Annotations are level 2/3 according to the 2007 metabolomics standards initiative (Sumner et al., 2007). Nodes **1-9**: Tomatidine molecular family. Molecular family **10**: a molecular family identified based on overlap of grape and fermented grape samples with plasma samples; multiple nodes contain spectral matches, however there is no library annotation and would otherwise remain completely uncharacterized. **d**. Summary of the spectra observed in plasma at each of the five food ontology levels. As this cohort received controlled diets, food categories observed in plasma samples were matched with the known foods consumed. Solid circles represent MS/MS matches to foods consumed during the study, while grey circles represent MS/MS matches to fermented versions of foods consumed, indicating possible byproducts of digestion. Open white circle indicates consumption was not recorded in this study.

A critical aspect of being able to leverage the food reference data, akin to expressed sequence tags, is that the associated metadata can be retrieved and organized. We leverage the Global FoodOmics ontology to identify different food categories in which MS/MS spectra are observed. These food counts can be summarized for a dataset and then visualized as a flow chart (**Figure 1d**). Due to the controlled research diets of the participants of the sleep and circadian study in **Figure 1d**, we were able to report if a given food category was consumed during the study. Of the 15 categories observed at level 5 of the food ontology, 8 represented direct matches, 3 represented fermented counterparts of consumed foods (such as yogurt and fermented grapes when milk and grapes were consumed), and 4 categories were not documented to be consumed, while coffee and tea were not provided to participants during this study. By and large, consistent with the lack of consumption of caffeinated beverages, evidence of coffee or tea consumption was only observed in two individuals. In one individual, caffeine was only detected in the first 48 hrs, and in the other volunteer, caffeine was observed in a single time point in the later part of the study (second to last time point). Spectral matches to caffeine were not detected in any of the other participants. Thus, the empirically-recovered food ontology information from metabolomics data demonstrates that these matches are consistent with the food that was consumed in this study.

To illustrate the broad utility of the Global FoodOmics reference data in enhancing the information gained from untargeted metabolomics, we co-analyzed the Global FoodOmics dataset with 27 human datasets (**Table S2**; at 1% FDR spectral matching), with the inclusion of additional study specific foods (SSF) where applicable (**Figure 1a**). These datasets contained between 5 and 2123 samples, represented multiple different biofluids and tissues, and included both adult and pediatric subjects, in conditions ranging from extremely long lived, such as a centenarian-enriched population in the Cilento Blue Zone in Italy, to inflammatory bowel disease, the healthy young adults undergoing experimental sleep restriction and circadian misalignment highlighted in **Figure 1** (Sprecher et al. 2019), children with medical complexity, adults with Alzheimer’s disease, and Covid-19 infections in Brazil (**Table S2**).

Spectral matching to food reference data, observed as overlaps between datasets from molecular networking, increased the interpretable fraction by 5.1 +/- 3.3 fold, even when compared to the library of all 150,633 public reference spectra that are used by the GNPS analysis infrastructure for annotation of public data which presently includes 29 spectral libraries, including from the three MassBanks (Japan, EU and North America) (Horai et al., 2010), HMDB (Wishart et al., 2018), ReSpect (Sawada et al., 2012), NIH natural product libraries (Huang et al., 2019), PNNL lipid library (Kyle et al., 2017), Bruker/Sumner, FDA libraries, Gates Malaria library, EMBL library, as well as many other GNPS contributed libraries (https://gnps.ucsd.edu/ProteoSAFe/libraries.jsp) and the commercial NIST17 library (CID portion only). Adding in additional information from molecular network connectivity, which can capture metabolized versions of molecules, the fold change of interpretable data increased further to 6.8 +/- 3.5 fold (**Figure 2**). The Global FoodOmics reference samples significantly increased the interpretation of various human metabolome samples above the initial annotation rate by 26.8+/- 3.3% for stool data (*P =* 2.8e-16, Games-Howell test), 27.5 +/- 5.2% for plasma data (*P =* 0.0040, Games-Howell test) and 41 +/- 4.6% for other human data (*P =* 0.00020, Games-Howell test). Further inclusion of connected nodes, representing potential metabolism via molecular transformations, results in a total increase of 43.7 +/- 3.1% (fecal; *P =* 6.9e-10, Games-Howell test), 51.2 +/- 6.9% (plasma; *P =* 2.8e-06, Games-Howell test), and 58.0 +/- 4.2% (human other; *P =* 1.4e-06, Games-Howell test) percent of MS/MS spectra that can now be leveraged as potentially a direct empirical readout of diet.

**Figure 2.**
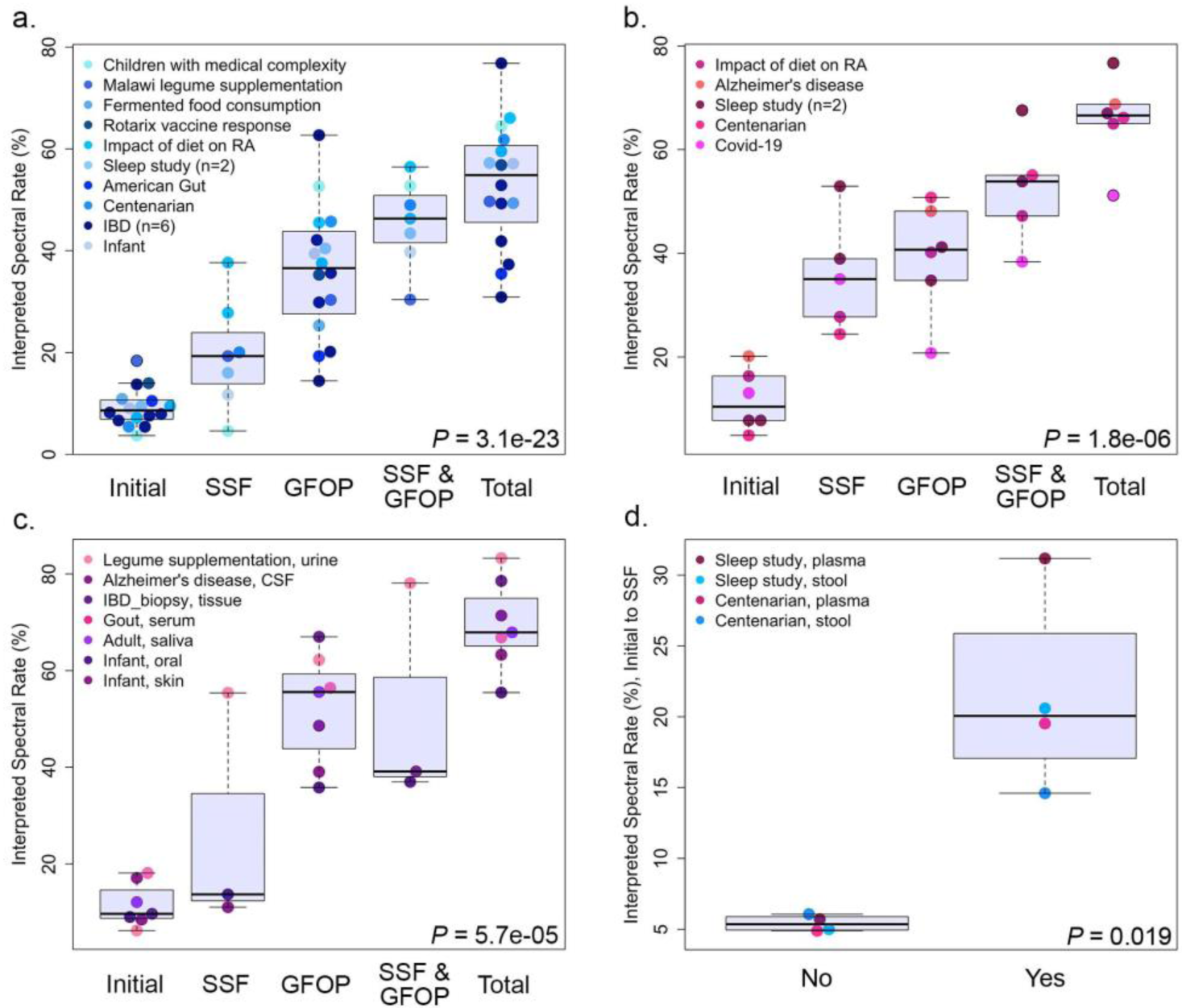
Increases of MS/MS spectral match rates when using interpretive metabolomics at the data set level. Spectral match rates of molecular features due to library match, food reference data, and molecular networking in **a**. stool data. Significant differences are determined by Welch’s F-Test. Library spectral matches (initial), spectral matches to study specific foods (SSF), spectral matches to Global FoodOmics project (GFOP) data, both (SSF & GFOP), expansion with molecular networking (Total). **b**. plasma data, and **c**. other human biospecimens. **d**. A crossover experiment between the centenarian data from Italy and the sleep and circadian study from the US, for both fecal and plasma samples. Study specific foods consumed by those individuals (yes) vs a different set of study specific foods (no), (Welch’s *t-*test).

For 14 of the public datasets, food samples of the region or exact dietary items frequently or exclusively eaten by that particular population were also collected (study specific foods; SSF). SSF and Global FoodOmics reference samples were separately (SSF; GFOP) and jointly (SSF & GFOP) evaluated for changes to the interpretable fraction of MS/MS spectra (**Figure 2**). For example, adding SSF (n=38) alone increased the percent of interpreted spectra for the centenarian stool data from an initial 5.4% annotation rate against spectral libraries to 20.0% interpreted (**Figure 2a**) and 4.9% initial to 24.4% for plasma samples (**Figure 2b**), and adding Global FoodOmics further expanded this to 49.0% (55.0% in plasma). For the sleep restriction and circadian misalignment study highlighted in **Figure 1**, the interpreted fraction also increased from an initial 7.2% to 27.8% (n=197 food samples; 45 of which are pooled meal samples), with a further increase to 46.3% when using the Global FoodOmics reference data set (7.8% to 38.9% and with Global FoodOmics up to 54% for plasma). Overall, the inclusion of SSF significantly contributed to the increase in dietary spectral matches in plasma (**Figure 2b;** *P =* 0.0028, Games-Howell test). In addition, in some cohorts the interpreted spectral rate reaches almost 80% after expansion with molecular networking (**Figure 2c**).

To further demonstrate that spectral matching using reference matching reflects dietary components, we performed a crossover study to test whether a mismatched SSF inventory would yield similar results to the increases observed across studies with SSF (e.g. centenarian foods for the sleep and circadian study cohort). Crossover revealed that the reciprocal tests interpretation rates were only a few percent (5–6%) in comparison to when the correct SSF were used (15–30%) (**Figure 2d**).

As the Global FoodOmics reference database expands with regionally-specific foods through a continued community effort, the interpreted fraction will likely increase. For example, when legume food data (15 files; SSF) similar to legumes supplemented in an infant malnutrition study were included in addition to the Global FoodOmics data, the number of spectral counts for legumes went from 105 to 2430 unique MS/MS spectra that matched, while other food categories such as dairy and meats remained constant (level 3 food ontology; Legume supplementation, urine). Regional specificity was also directly evident for plasma samples collected in Brazil for a Covid-19 study, which displayed more spectral matches to a locally collected set of 60 Brazilian food samples with ∼35% increase than to the entire Global FoodOmics reference dataset, that is dominated by US food, which only gave an ∼20% increase in spectral matches (**Figure 2b**). Thus, although there is some overlap among the data from different foods, and even overlap among human-derived metabolites and the food data (e.g. many primary metabolites or those common in vertebrates), a large proportion are sufficiently unique to reveal, at least in part, the dietary composition in the study.

To assess if interpretive metabolomics could be used to empirically establish adherence to dietary recommendations using MS/MS data, we analyzed a data set from rheumatoid arthritis patients (RA) asked to follow an anti-inflammatory diet (ITIS diet) (Bustamante et al., 2020). We compared the per sample extracted food counts with the recommended diet alteration as well as self-reported diet diary entries. The recommended diet included some foods to be avoided (such as coffee, refined sugars and milk), some foods to be restricted (minimize red meat and egg consumption) and some foods to be frequently consumed (such as fruits/vegetables, and plain unsweetened yogurt). In total, 47 foods and beverages were observed in this project with interpretive metabolomics (**Figure 3a**). By and large, most adhered to the recommended diet, as food counts of recommended foods increased, and those of foods to avoid decreased. Although there are instances when the mass spectrometry based observations did not match the recommended diet regime, the self-reported dietary records matched the empirically determined foods better than the recommended dietary information (**Figure 3b**). We further validated these matches using source tracking with 16S rRNA gene amplicon data collected on ∼1500 samples of the Global FoodOmics foods, to predict food source contribution to the RA study stool samples. We observed a highly significant correlation in the proportion change of food sources predicted in the stool samples and metabolites in the plasma before and after dietary intervention (Pearson r = 0.57, p-value = 0.003; **Figure S2**). The empirically recovered food ontology information from interpretive metabolomics, in conjunction with validation with DNA sequence data, demonstrates the ability to recapitulate dietary readouts from human biospecimens and assess diet adherence.

Interpretive metabolomics comes with several caveats to consider. We are not yet able to capture a complete picture of the human diet: for example, in the RA study, the participant diet diaries contained foods not yet captured in the FoodOmics database, potentially leading to an underestimation of food types observed. Community expansion of the Global FoodOmics database with specific foods and food ingredients will ultimately eliminate this issue.

**Figure 3.**
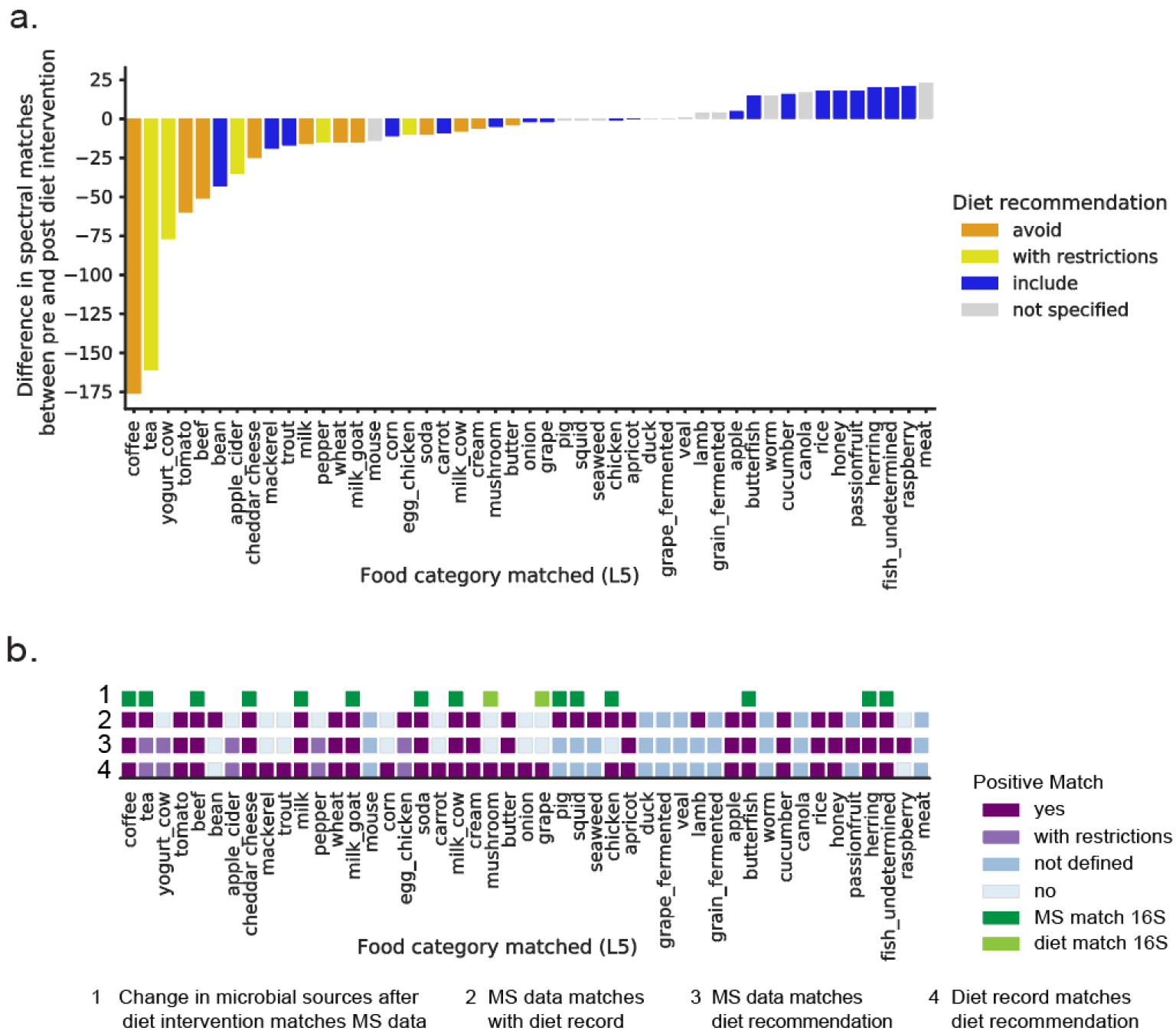
Using interpretive metabolomics to assess dietary recommendations at the study level. **a**. Food ontologies of a rheumatoid arthritis cohort before and after a specific dietary recommendation of a low inflammatory diet. Plasma data are used. Food categories indicated as ‘with restrictions’ encompass foods where different types are encouraged and others discouraged (green vs. black tea) or foods that were supposed to be minimized (such as limiting egg consumption to 2 eggs per week). Food categories indicated as ‘not specified’ could not be matched to the suggested diet. **b**. Comparison of interpretive metabolomics results in the recommended diet and self-reported diet intake. Diet diaries were tabulated as consumption or no consumption of >200 food categories over the 28 days of the study and matched to the MS food categories, as possible. Matches to 16S rRNA gene sequence data are based on Bayesian source tracking proportions from the bacterial community, with food types as sources and rheumatoid arthritis fecal samples as sinks. The increase or decrease in the proportion of food source contribution pre and post dietary intervention (y-axis) is colored according to dietary recommendations.

Another consideration is similar to what is observed with expressed sequence tags/RNASeq, where it is common to observe that there are multiple sample types, tissue locations or conditions that result in misinterpretation because the same sequence occurs in multiple locations. By analogy, a molecule could be produced by humans but also be part of different diet sources (i.e. cholesterol produced by the human body versus consumed). However, such matches still enable one to formulate a hypothesis that the observed MS/MS features from the human data might originate from the reference data as source, in this case food, especially when there are hundreds or thousands of signatures that point to specific foods or food groups that overlap.

As we saw in many of the above datasets, it is not atypical to observe small numbers of spectral matches to insects, rodents, fungi and worms within diet read-outs. Although data on fungi, tarantula, crickets, and black ants, meant for human consumption, are included, most of these samples that match human data sets are from a Global FoodOmics sampling effort at the San Diego Zoo. While there is likely some overlap with molecules from these less common foods to those that humans more commonly consume (e.g. certain acylcarnitines might be found in beef and mice), the FDA food contamination guidelines allow for insect, fungal, worm, rodent parts and fecal matter to be present in food in quantities that surprise many non-specialists (Center for Food Safety and Nutrition, 2019) For example, peanut butter is allowed to have 30 or more insect fragments and one rodent hair per 100 grams, and apple butter is allowed to have “5 or more whole or equivalent insects (not counting mites, aphids, thrips, or scale insects) per 100 grams of apple butter.” As long as these dietary “additives” are added to the reference data set, they too will be observed. Thus, interpretive metabolomics can provide empirical support for dietary compliance in nutritional content, including in clinical studies, and capture information that would otherwise remain hidden.

## Conclusion

Here we show that well-curated reference datasets can be leveraged to provide a deeper understanding of untargeted metabolomics. Adding food-based spectral matches improves our ability to interpret molecular features 2 to 14-fold, and further improves to 3 to 17-fold by incorporating connections from molecular networking, providing a deeper insight of the metabolomic “dark matter.” Our results indicate that a direct empirical readout of diet adherence is within our reach using interpretive metabolomics, by combining structural, source, and chemical similarity measures.

Although we demonstrated the power of interpretive metabolomics with food data as reference, any individual reference data set or combination of multiple data sets could be used in this fashion. We envision the broad application of such an approach. Generating databases for environmental allergens, medications, illegal substances, food ingredients and personal care products can inform within those research areas on potential exposures and food adulteration. Further, such investigations may also have far reaching impacts to understand commonalities that underlie different diseases. Over time, as metabolomics data repositories begin to control metadata vocabularies, most public data could be leveraged and reused as a reference data set on its own. This will significantly improve the interpretability of all metabolomics data, be it from environmental, animal, or human sources.

## Acknowledgments

Funding sources: We thanks the CCF foundation #675191, U19 AG063744 01, R01AG061066, 1 DP1 AT010885, P30 DK120515 Office of Naval Research MURI grant N00014-15-1-2809 and NIH/NCATS Colorado CTSA Grant UL1TR002535. This work was also supported in part by the Chancellor’s Initiative in the Microbiome and Microbial Sciences and by Illumina, Inc. through reagent donation in partnership with the Center for Microbiome Innovation at UC San Diego. We would like to thank Elaine Wolfe and Karenina Sanders for sample processing, and Jeff DeReus for data handling, processing and maintaining the computational infrastructure. JPS was supported by SD IRACDA (5K12GM068524-17), and in part by USDA-NIFA (2019-67013-29137) and the Einstein Institute GOLD project (R01MD011389). RC and MG were supported by Krupp Endowed Fund. RHM was supported through a UCSD training grant from the NIH/NIDDK Gastroenterology Training Program (T32 DK007202). The Brazilian National Council for Scientific and Technological Development (CNPq)-Brazil [245954/2012] to MFO. KS was supported by a PROMOS fund (DAAD). WB is a postdoctoral researcher of the Research Foundation – Flanders (FWO). We thank Ricardo da Silva for his feedback and early bioinformatics analysis for the Global FoodOmics project. We further acknowledge all the individuals that contributed samples as well as companies and organizations that have donated samples: Townshend’s Tea Company, BDK Kombucha, Oregonian Tonic, Squirrel & Crow, Venissimo cheese, Fermenter’s Club San Diego, Good Neighbor Gardens, Sprouts Farmers Market, Ralphs, Whole Foods and SD Zoo and Safari Park. Specifically thank you to Austin Durant for coordinating sampling at Fermentation Festivals and the wonderful staff at SD Zoo Global for coordinating and helping with sample collection: Michele Gaffney, Edith Galindo, Katie Kerr, Andrea Fidgett, Jennifer Stuart, Debbie Tanciatco, and Lisa Pospychala.

## Author Contributions

PCD, RK, RJD, and JMG conceptualized the idea.

MWP, FDO, KCW, CMA, EB, KS, PCD, RJD, RK, NCS, ADS, GA, DM, NPL, and JMG collected

FoodOmics samples and performed metadata curation.

MWP, FDO, FV, CMA, EB, NCS, and JMG performed FoodOmics sample processing and MS data acquisition.

AJJ, PBF, ED, QZ, DN, DM, JPS, and JMG curated Global FoodOmics metadata to match FNDDS.

JBW, BSB, BJB, RC, MGDB, MD, EOE, DG, LH, JK, MM, CM, RK, KES, DVR, CW, KPW, MFO, RHM, DW, RT, JGA, PD, MG, DG, AKJ, BJB, RMS, KCW, ADS, FV, NPL, and JMG provided samples, comparative dataset, and/or detailed metadata.

LMMM, TMC performed Covid-19 patient and/or food sample preparation and analysis. PLJ was the physician responsible for the Covid-19 patients (provided samples).

RDRO was the physician responsible for collecting the plasma from Covid-19 patients (provided samples).

FPV was responsible for tabulation of Covid-19 patient data and analysis of health components, and body dimensions.

TS, MB, LDG, GH performed 16S sequencing and prep.

CM, DM, JPS performed source tracking and/or 16S data analysis.

MW supported GNPS computational infrastructure used in the study.

CLW, WB, AKJ, ES, AT, NPL and JMG analyzed MS data.

CLW, WB, AKJ, CM, and JMG generated figures.

PCD, RK, RJD, ADS, and JMG supervised the work.

PCD, RK, CLW, and JMG wrote the paper.

All authors have contributed feedback and edits to the manuscript.

## STAR Methods

### Resource Availability

#### Lead Contact

Further information and requests for resources should be directed to and will be fulfilled by the Lead Contact, Pieter Dorrestein.

#### Materials Availability

This study did not generate new unique reagents.

#### Data and Code Availability

The code generated during this study is available on GitHub at https://github.com/DorresteinLaboratory/GlobalFoodomics.

Raw and processed 16S rRNA amplicon sequencing data is available at Qiita study #11442 and raw sequence data has been deposited at EBI accession ERP122648.

GNPS task ID of analysis used for tag generation: f1a1f3a61aca416a9b3687d72488da7f The following files are available in addition to the Global FoodOmics mzXML files on massive.ucsd.edu under MSV000084900: metadata as a .txt; an image repository with between 1 and 6 images per food; table of FDR-based parameters; raw food count data for RA study; full size PDF of sleep restriction and circadian misalignment study - GFOP3500 molecular network (excerpts found in Figure 1).

Metadata dictionary: https://docs.google.com/spreadsheets/d/1Ebn-TgMWEkd_7KOw9TCRvHGPsE7dGjVCr7dg28pwbmM/edit#gid=727944641

The GNPS analyses used in this study can be accessed on-line at the following links:

- Sleep study (MSV000083759; https://gnps.ucsd.edu/ProteoSAFe/status.jsp?task=e0bf255bcb2e492bb0be3be1a691b5fb; https://gnps.ucsd.edu/ProteoSAFe/status.jsp?task=6fe434761daf4f9da540cf1fd90b3985; https://gnps.ucsd.edu/ProteoSAFe/status.jsp?task=9a90bd12f51e453e968656e6458e0da4)
- Centenarian (MSV000084591; https://gnps.ucsd.edu/ProteoSAFe/status.jsp?task=8895b6e3445546c4a5bc3a726a920227; https://gnps.ucsd.edu/ProteoSAFe/status.jsp?task=981c9a7d39f742bda296d52f856981e5)
- Impact of diet on RA (MSV000084556; https://gnps.ucsd.edu/ProteoSAFe/status.jsp?task=0794151fce2c4c18a7a0aa3a09140169)
- LP Infant (MSV000083462; MSV000083463; https://gnps.ucsd.edu/ProteoSAFe/status.jsp?task=a7b222466ef844e69cdbd9835d2f6c39; https://gnps.ucsd.edu/ProteoSAFe/status.jsp?task=c756a9dfb5c34a2a8655f88114edf0a8; https://gnps.ucsd.edu/ProteoSAFe/status.jsp?task=4a322e640bb644068030949267fb4ea9)
- Children with Medical Complexity (MSV000084610; https://gnps.ucsd.edu/ProteoSAFe/status.jsp?task=df24423835a341969342c2086b46275a)
- American Gut (MSV000081981; https://gnps.ucsd.edu/ProteoSAFe/status.jsp?task=4884483bcffe4f269819858c3fd4faef)
- Fermented food consumption (MSV000081171; https://gnps.ucsd.edu/ProteoSAFe/status.jsp?task=5cca39e0ebab4066a56e41ded48b4466)
- Malawi legume supplement (MSV000081486; https://gnps.ucsd.edu/ProteoSAFe/status.jsp?task=93ba727aa9234727a73ae7860b2af3ca)
- Rotarix vaccine response (MSV000084218; https://gnps.ucsd.edu/ProteoSAFe/status.jsp?task=08e9b9e048f04ac4b416e574a073e8e6)
- IBD_1 (MSV000082431; https://gnps.ucsd.edu/ProteoSAFe/status.jsp?task=ec08eed8f186430d893c63111409baf4)
- IBD_individual (MSV000079115; https://gnps.ucsd.edu/ProteoSAFe/status.jsp?task=fad746939afd4184975a296436aebfb7)
- IBD_seed (MSV000082221; https://gnps.ucsd.edu/ProteoSAFe/status.jsp?task=907f2e0b7878417dbdb4c83f0df0e83a)
- IBD_biobank (MSV000079777; https://gnps.ucsd.edu/ProteoSAFe/status.jsp?task=a79fbd4c96124209adfd0ef84cb56dec)
- IBD_2 (MSV000084775; https://gnps.ucsd.edu/ProteoSAFe/status.jsp?task=07f855658c5342458045032ea70fc526)
- IBD_200 (MSV000084908; https://gnps.ucsd.edu/ProteoSAFe/status.jsp?task=55bef02250d744eb97c6040c379cbfb4)
- Alzheimer’s disease (MSV000085256; https://gnps.ucsd.edu/ProteoSAFe/status.jsp?task=aac78e9d23b84194ab2f768cb685c636)
- Covid-19 (MSV000085505; MSV000085537; https://gnps.ucsd.edu/ProteoSAFe/status.jsp?task=9cbcb6b46fe24826bc56c9e893d0bd2b)
- IBD_biopsy (MSV000082220; https://gnps.ucsd.edu/ProteoSAFe/status.jsp?task=a83a279dad154f9ca7b549d40ce117ba)
- Gout (MSV000084908; https://gnps.ucsd.edu/ProteoSAFe/status.jsp?task=55bef02250d744eb97c6040c379cbfb4)
- Adult Saliva (MSV000083049; https://gnps.ucsd.edu/ProteoSAFe/status.jsp?task=6dd6e5b1cf454d67b8a2b3c151c18f4a)
- Legume supplementation (MSV000084663; https://gnps.ucsd.edu/ProteoSAFe/status.jsp?task=93ba727aa9234727a73ae7860b2af3ca)

Networking parameters were set based on the MOLECULAR-LIBRARYSEARCH-FDR workflow on GNPS with the following task IDs:

- GFOP3500: a7bf6cc3f91d466bab923f2268d6f4fc
- Sleep deprivation: b55ab4004ed342d7b4ed1c488e935998
- Sleep study: 78bbfed8574748d1a77dc7c2f1a44d39
- Sleep study_SSF_test: b55ab4004ed342d7b4ed1c488e935998
- Centenarian: 265a9553c69e47499cca3de056b43178
- Centenarian_SSF_test: 265a9553c69e47499cca3de056b43178
- American Gut: aee5dde3b2f84079a264e68ec981487e
- Fermented food consumption: a44d1b2e1b9d4612974d0b85021675a7
- Malawi legume supplement: de7b55f8adaa4ad9b2a8430e30435bf3
- Children with Medical Complexity: f27243af071b43ab90d846bda959fc1c
- Rotarix vaccine response: a2e02e3f97a54ca08e3866cc60f8d42b
- Impact of diet on RA: 62b8754e761549f3b94ffae83d7ab95a
- LP infant: 532aba2ad3644fadba0e6e7ea063c7ee
- IBD_1: bb10b1ce90a24f3a9cef1e85e88c3882
- IBD_biopsy: c4cfda90933b4842a7154f5f2def139d
- IBD_individual: 3ce8cc636ae944848b4ada322aaf12fe
- IBD_seed: ebbb715fc605457ba5f7e910b79d6177
- IBD_biobank: 9465c34cf5444e12b89318b1fb363714
- IBD_2: 983fa9271136404fb5743b44a6a109f0
- IBD_200: e5acf5726722486caa897b2b07d402e8
- Impact of diet on RA: 62b8754e761549f3b94ffae83d7ab95a
- Alzheimer’s disease: 658103164325425981c097cecba840b0
- Gout: a478f419ae824378aa02e5e1b310cad2
- Adulta saliva: 32980f95dbd5437aaa9e15d05c7246bb
- LP infant: 8bfbdc1bf38c418fb223306cd42af897
- LP infant: 3e414e13a4394bb78c07f7ca7f4d1be3
- Legume supplementation: 2ca007303b9c4bb3820f392b996eba27
- Alzheimer’s disease: 658103164325425981c097cecba840b0
- Covid-19 Brazil: d16eb32276c84bdb9c35c5872e97a986

## Methods

### IRB information for the human datasets used in this study, made public on MassIVE

Sleep study (MSV000083759; IRB 15-0282), centenarian (MSV000084591; IRB 180478), Impact of diet on RA (MSV000084556; IRB 161474), LP Infant (MSV000083462; MSV000083463; IRB 151713 UCSD), Children with Medical Complexity (MSV000084610; IRB 161948 UCSD), American Gut (MSV000081981; IRB 141853 UCSD), Fermented food consumption (MSV000081171; IRB 141853 UCSD / published), Malawi legume supplement (MSV000081486; IRB ID #201503171; Washington University Human Studies Committee), Rotarix vaccine response (MSV000084218; IRB is PR-10060 from University of Virginia), IBD_1 (MSV000082431; IRB # 150675), IBD_individual (MSV000079115; IRB # 150675), IBD_seed (MSV000082221; UCSD HRRP 131487), IBD_biobank (MSV000079777; UCSD HRRP 131487); IBD_2 (MSV000084775; IRB # 150675), IBD_200 (MSV000084908; IRB # 150675), Alzheimer’s disease (MSV000085256; UCSD IRB # 170957), Covid-19 (MSV000085505; MSV000085537; IRB approval number is 30248420.9.0000.5440 (University of São Paulo, Brazil), IBD_biopsy (MSV000082220; IRB number is 120025), Gout (MSV000084908; IRB Project #160768X), Adult Saliva (MSV000083049; IRB 150275 UCSD), Legume supplementation (MSV000084663; IRB ID #201905103).

### Global FoodOmics

#### Sample Collection

Sampling methodology was developed in order to facilitate sample collection in any environment, from the home, a restaurant, a festival, or in the lab. Initial samples were collected in a consistent manner, between April 2017 and March 2018. Additional sets of samples were added through Fall 2019. Each sample was assigned a unique number identifier upon sampling, which was used to trace the origin of the sample, and to organize descriptive information about the sample. In addition, when possible samples were photographed by the participant to create a photographic archive of all samples (uploaded to MassIVE MSV000084900; >4000 images representing 67% of the samples (2399/3579)). Primarily for the initial data set these images were used as the first point of reference for the collection of ancillary information about the different samples (termed metadata, described in more detail below). The image archive was critical, because as the project evolved and the breadth of sample types increased, new categories were added to the metadata, which were then filled in weeks or even months after sample collection.

Samples were frozen at −80°C within 24 h of sample collection, unless otherwise noted in the metadata. Two samples were collected for each food or beverage included in the study. One sample was collected as an archive and directly frozen, and a second sample was collected for extraction. Food samples were collected in a tube prefilled with 1 ml 95% ethanol (Ethyl alcohol (Sigma-Aldrich) and Invitrogen UltraPureTM Distilled Water), as high ethanol concentrations are efficacious at preserving the sample for both DNA and metabolite analyses (Song et al., 2016). Samples were collected into 2 ml round bottom microcentrifuge tubes (Qiagen) and weighed prior to freezing. The pre-sample and post-sample weights as well as the weight differences were recorded in the metadata. It was not possible to collect all samples at a given concentration of extraction solvent (ethanol), because sampling was performed in many different environments and is meant to be consistent with future crowd-based community science participation. Therefore, the data can be compared qualitatively and not quantitatively, however for certain subsets 50 mg of material was collected consistently.

Additional sets of food samples were added to the core set using the same methods as outlined above when possible. Samples from Venezuela were collected whole in absolute ethanol >=99.8% (Sigma Aldrich) and the extract was processed directly.

The experimental protocol for the sleep restriction and circadian misalignment study has been described previously (Sprecher et al., 2019). Meals and food samples were prepared by the Clinical and Translational Research Center Nutrition Core of the Colorado Clinical and Translational Sciences Institute. Food was transported to the research site and refrigerated for the duration of the in-patient study. Individual meals were sampled and stored frozen in ziploc bags. They were stored at −80° C prior to subsampling and LC-MS/MS analysis. Images are contained in a separate Sleep Study folder (MSV000084900).

For several of the human studies we collected data on associated foods, which were processed according to the same methods as the Global FoodOmics samples. The number of SSF samples per cohort are outlined here: experimental sleep deprivation (197 samples; 45 are pooled); centenarian (38 individual samples); malawi legume supplement (14; 2 sample types, several extraction types); children with medical complexity (24 formula samples; 11 exact overlap); RA diet samples (20 individual sample; 2 samples types (stool, plasma), 2 time points)); mother’s milk (58 milk samples); legume supplements (15 individual legume samples; 6 different types).

#### Community-based science collection

The first sample collected was a carrot from a home garden. The participant was interested in how the soil conditions from prior tenants would impact the chemistry of the carrot, since the gardening practices of the prior tenant were unknown (organic or not, pesticide usage, etc.). In addition, home grown foods often taste different than store bought, likely reflected in the food metabolome.

During the course of sampling, samples were received from over 50 different individuals in California as well as from different states as well as countries (such as Venezuela, Italy and most recently Brazil). Contributions from individuals ranged from produce from home gardens, home fermented products (yogurt, kombucha, sauerkraut), meat and dairy from private farms, to items individuals had purchased that were of interest to them.

We were also directly invited to sample at local stores and organizations, including Venissimo cheese, Good Neighbor Gardens, and the San Diego Zoo and San Diego Zoo Safari Park, as well as local supermarkets such as Sprouts Farmers Market, Whole Foods Market, and Ralphs. We were invited by San Diego Fermenter’s Club founder Austin Durant to the San Diego Fermenter’s Club meeting and sampled from multiple vendors at both the Oregon Fermentation Festival in 2017 as well as the San Diego Fermentation Festival in 2018. We also received citrus samples from a farm at the US-Mexico border, with visibly dark skin due to air pollution, a particular concern of the farmer. Other sampling occurred in conjunction with study design, as was the case for the Rheumatoid arthritis cohort and the Covid-19 study. In total we engaged with a broad range of individuals, organizations, businesses and scientists, to generate this dataset of 3579 samples (for future use this is already expanded beyond this number due to the collection of sets of SSF). A predominance of foods included in this initial dataset were sampled and/or purchased in California, leaving room for much further expansion and the inclusion of a crowd-sourced community science initiative to expand the array of samples.

The sample set contains a broad set of simple foods including fruits, vegetables, grains, as well as raw meat and fish, which build the foundation of many food products. In addition, we have 1133 fermented samples. This subcategorization of foods is made possible by the metadata collected on these samples, described in the Metadata Curation section. The breadth of samples included in the dataset necessitated careful collation and a range of information about the samples, resulting in 157 different metadata categories to describe various aspects of these food and beverage samples.

Samples originate from over 50 different identified countries of origin (Argentina, Australia, Austria, Belgium, Bolivia, Brazil, Canada, Chile, China, Columbia, Croatia, Ecuador, England, Ethiopia, France, Germany, Greece, Guatemala, Haiti, Holland/Netherlands, India, Indonesia, Ireland, Israel, Italy/Sardinia, Japan, Kenya, Korea, Madagascar, Malawi, Mexico, New Zealand, Nilgiri, Peru, Philippines, Poland, Serbia, Portugal, Russia, Scotland, South Africa, Spain, Switzerland, Taiwan, Thailand, Trinidad & Tobago, Turkey, UK, USA/Puerto Rico, Vietnam, Venezuela; EU, South America not included separately).

#### Metadata Curation

##### General organization

Detailed information about each sample was captured in the form of metadata. The metadata are in the form of an array, where each row represents one sample and each column captures unique information about the sample (See Supplementary Information for Metadata File, as well as updates on Massive MSV000084900). This matrix allows for the categorization of foods by various different attributes and links these attributes to the sample numbers, the data files (.mzXML filename), as well as the 16S sequence information on Qiita (sample_name). The initial metadata categories captured included sample description, sample number, location sample was collected, the weight of the sample (pre-sample, post-sample, sample weight), the day it was collected, and whether an image had been taken and renamed to match the sample number and archived in the image repository. The initial 9 categories captured minimal information and allowed tracking of information about the sample.

During the process of sample collection, the diversity of the samples being collected necessitated the addition of columns to capture more information about the samples and to be able to categorize them and compare different attributes. These columns grew to capture highly detailed information about each sample, for example whether the sample was organic, if it was raw or cooked, if it was washed before sampling, or for cheese samples whether it is the rind or the curd, etc. As columns were added, the initial columns and the image repository were used to trace back information.

##### Classification scheme

Various classifiers are used to describe foods, however we were unable to find an established scheme able to capture the diversity of samples, as well as distill the metadata down into a manageable number of categories to distinguish differences between the metabolomes of different food classes. We therefore categorized the foods by sample_type, which captured whether the sample was a food, beverage, or other item (for example supplements) and then expanded and shaped a unique categorization which takes into account the species and botanical definitions of foods. The sample_type categories range from sample_type_land_aquatic, to differentiate items sourced from different physical environments, sample_type_common, which allows for representation of a particular food group which was not otherwise captured in the metadata, such as zoo food or candy. The sample_type groups also include a hierarchy from group1 to group6 (Levels 1 through 5 are referenced in this manuscript), specific to foods and groupB1 through groupB3 which contain beverage specific information (alcoholic [binary], carbonated [binary], type of beverage [such as red wine, kefir, soda, etc.]).

##### Complex samples

The above classification scheme gave sufficiently detailed information about simple foods (ones that have only one ingredient and could thus be filled out to the last group level, such as red cherry tomato). Complex foods contain not only multiple ingredients, but include highly processed foods purchased with ingredient lists as well as home cooked or restaurant meals. These foods have a higher variability of information known about them. The top 6 ingredients are captured in individual metadata categories, with a seventh ingredient field which contains the remainder of the ingredients (if known). However, the order of ingredients does not always clearly reflect the type of food and some constituents that may be of interest, such as tree nuts which may only be found in trace quantities. The sample_type_common category captured some of the information about the type of sample (candy), however to have a tangible classification of different ingredient types, we generated a specific complex food ontology based on the known presence of common categories (corn, dairy*, egg*, fruit, fungi, fish*, shellfish*, meat, peanut*, seaweed, soy*, tree nut*, vegetable/herb, wheat* (*designates known food allergen)). These categories reflect the main food groups and some of the most common allergens (US FDA Food Allergen Labeling And Consumer Protection Act of 2004) (Sicherer and Sampson, 2006), items which are of interest when correlating food metabolome data with other datasets, such as human fecal material (where the foods eaten are known or unknown).

##### Fermented foods

Preservation and processing method are included in the metadata. However, due to the potential importance of fermentation in the alteration of the food metabolome, and the potential health benefits that have been ascribed to fermented foods, several categories were included to highlight this feature: fermented or not, whether it contains live active cultures, whether it contains chocolate (which then was cross checked with the fermented category, as chocolate is a fermented food). The list of fermented foods crosses many of our sample types as it includes fermented dairy (yogurt, cheese), fermented meat/fish (salami, fish sauce), fermented vegetables (kimchi, sauerkraut), fermented fruit (chocolate, coffee), and fermented grains/legumes (bread, tempeh).

##### Food specific categories

Certain individual food categories also necessitated creation of specific categorization. For example, cheeses have the specific categories cheese_part (curd vs. rind), cheese_type (washed, blue, etc), and cheese_texture (soft, semi-soft, semi-hard, hard). Particularly for raw plant products, such as fruits, vegetables, grains which form the basis for many food ingredients, we captured botanical information: botanical_anatomy (fruit, leaf, tuber, seed, etc.), botanical_genus, and botanical_genus_species (when known). Tea samples have tea quality and tea type as distinct categories.

#### Metadata for Cross-study Comparison

To facilitate cross study comparison, we included the Earth Microbiome Project ontology: empo_1 (level 1: Free-living, Host-associated, Control, or Unknown), empo_2 (level 2: Saline, Non-saline, Animal, Plant, or Fungus), and empo_3 (level 3: most specific habitat name) [http://www.earthmicrobiome.org/protocols-and-standards/empo/]. Wherever possible we linked foods to food identifiers or created identifiers and categories that built upon the existing framework as defined by the U.S. Department of Agriculture’s Food and Nutrient Database for Dietary Studies 2011-2012 (FNDDS) food grouping scheme (Martin et al., 2012).

#### Sample Preparation

A sterile stainless steel bead was added to each sample collected in ethanol and the samples were thawed on ice for 30 min. Samples were homogenized at 25–30 Hz for 5 min using a tissue homogenizer (QIAGEN TissueLyzer II, Hilden, Germany). Samples were swabbed with sterile dual tip swabs (BD swubes) and frozen immediately at −80°C until DNA extraction.

#### DNA Extraction and 16S rRNA gene amplicon sequencing

DNA extraction and 16S rRNA gene amplicon sequencing were performed using Earth Microbiome Project (EMP) standard protocols (http://www.earthmicrobiome.org/protocols-and-standards/16s) (Thompson et al., 2017). DNA was extracted with the Qiagen MagAttract PowerSoil DNA kit as previously described (Marotz et al 2017). Amplicon PCR was performed on the V4 region of the 16S rRNA gene using the primer pair 515f-806r with Golay error-correcting barcodes on the reverse primer. Amplicons were barcoded and pooled in equal concentrations for sequencing. The amplicon pool was purified with the MO BIO UltraClean PCR cleanup kit and sequenced on the Illumina MiSeq sequencing platform. Raw sequence data were uploaded to Qiita for pre-analysis processing (Qiita study ID: 11442) (Gonzalez et al., 2018). In Qiita, raw sequence data were demultiplexed and minimally quality-filtered using the QIIME 1.9.1 script split_libraries_fastq.py, with a Phred quality threshold of 3, allowing for reverse complemented barcodes and mapping barcodes, and default parameters. Demultiplexed, quality-filtered sequence data were then trimmed to a read length of 150-bp, denoised with Deblur v1.1.0 (Amir et al., 2017) using default parameters, and subject to fragment insertion with SATéEnabled Phylogenetic Placement (Janssen et al., 2018) into the GreenGenes 13.8 reference phylogeny (McDonald et al., 2012), using default parameters, to generate an inclusive phylogeny. An observation table of per-sample counts of Deblur sub-operational taxonomic units were output into BIOM format for analyses (n = 1511 samples). Outside of Qiita, we assigned taxonomy to denoised reads using QIIME2’s feature-classifier, classify-sklearn, using the GreenGenes 13.8 pre-fitted sklearn-based classifier (i.e., 99% OTUs, 515f/806r region of sequences), and default parameters (Bokulich et al., 2018; Bolyen et al., 2019).

#### SourceTracker analyses

SourceTracker 2.0.1 (http://github.com/biota/sourcetracker2) was used to predict the proportion of microbial source environment contributions to a sink using a Bayesian classification model together with Gibbs sampling (Knights et al., 2011). The Deblur 150-bp observation table consisting of 1511 food samples was used as the set of source environments and the Rheumatoid Arthritis (RA) data set consisting of 49 fecal samples was used as the sink. All source and sink samples were rarefied to 2000 sequences per sample before source-tracking and doubleton ASVs were removed. Leave-one-out cross-validation was used to predict the source samples with heterogeneity from all other food categories. After source sample filtering a total of 346 samples representing a total of 25 broad food categories were retained. Food microbial source contributions were then predicted for RA samples and the difference in food contribution before and after diet intervention was calculated and compared by diet recommendations.

#### Metabolite Extraction

Homogenized samples were incubated for 40 min at −20°C and centrifuged (Eppendorf centrifuge 5418, Hamburg, Germany) at 20,000 rpm for 15 min at 4°C. 400 μL of supernatant were transferred to a 96-well deep well plate and dried by centrifugal evaporation (Labconco Acid-Resistant Centrivap Concentrator, Missouri, USA). Dried extracts were reconstituted in 150 μL of resuspension solution (50% methanol with 2 μM sulfadimethoxine), then vortexed for 2 min and sonicated for 5 min in a bath water (Branson 5510, Connecticut, USA). Resuspended extracts were then centrifuged for 15 min at 20,000 rpm and 4°C (Thermo SORVALL LEGEND RT, Germany) and transferred to a 96-well shallow well plate, and diluted either 5x or 10x to avoid saturating the MS detector.

#### Liquid Chromatography - Mass Spectrometry

Food extracts were analyzed using a UltiMate 3000 UHPLC system (Thermo Scientific, Waltham, Ma) equipped with a reverse phase C18 column, prepended with a guard cartridge (Kinetex, 100 × 2.1 mm, 1.7 μm particles size, 100 Å pore size; Phenomenex, Torrance, CA, USA), at a column compartment temperature of 40°C. Samples were chromatographically separated with a constant flow rate of 0.5 ml / min using the following gradient: 1.5 min isocratic at 5% B, up to 100% B in 8 min, 3 min isocratic at 100% B, back to 5% B in 0.5min and then 1.5min isocratic at 5% B (A: H2O + 0.1% formic acid; B: Acetonitrile (ACN) + 0.1% formic acid (LC-MS grade solvents, Fisher Chemical, Hampton, United States)).

The UHPLC system was coupled to a Maxis Q-TOF Impact II mass spectrometer (Bruker Daltonics, Bremen, Germany) equipped with an electrospray ionization source. MS spectra were acquired in positive ionization mode using Data Dependent Acquisition (DDA) with a mass range of *m/z* 50–1500. The instrument was externally calibrated two times per day to 1.0 ppm mass accuracy using ESI-L Low Concentration Tuning Mix (Agilent Technologies, Waldbronn, Germany). Hexakis (m/z 622.029509; (1H, 1H, 2H difluoroethoxy)phosphazene (Synquest Laboratories, Alachua, FL)) was used for lock mass correction. MS/MS spectra were acquired for the top 5 ions in each MS1 spectrum, with active exclusion after 2 spectra (maintained for 30 seconds). Known contaminants as well as lock mass values commonly used with this instrument were added to an exclusion list (*m/z* values listed): 144.49–145.49; 621.00–624.10; 643.80– 646.00; 659.78–662.00; 921.0–925.00; 943.80–946.00; 959.80–962.00.

Raw high resolution mass spectrometry data files were converted to open source .mzXML format using Bruker DataAnalysis software after lock mass correction (*m/z* 622.0290). Raw data files as well as converted .mzXML files were uploaded to MassIVE (publicly available under unique identifier MSV000084900) and further analyzed on Global Natural Product Social Molecular Networking (GNPS) (https://gnps.ucsd.edu), as described below.

#### MS2 Data Processing

##### FDR estimation

False discovery rate (FDR) estimation was calculated using Passatutto analysis workflow in GNPS (Scheubert et al. 2017; Wang et al. 2016). FDR estimation was used to determine the cosine value required with a minimum of 5 matched peaks to achieve an FDR of 1%. See the Data availability section for accession information.

*Molecular networking using GNPS:* Molecular networking analysis and library search were performed using GNPS classical molecular networking release_18 (Wang et al. 2016). 3579.mzXML data files (available at MassIVE ID MSV000084900) were included in the analysis. The data were filtered by removing all MS/MS peaks within +/- 17 *m/z* of the precursor *m/z*. MS/MS spectra were window filtered by choosing only the top 5 peaks in the +/- 50 *m/z* window throughout the spectrum. The data was then clustered with MS-Cluster with a parent mass tolerance of 0.02 *m/z* and an MS/MS fragment ion tolerance of 0.02 *m/z* to create consensus spectra. Further, consensus spectra that contained less than 2 spectra were discarded. A network was then created where edges where filtered to have a cosine score above 0.65 (slight variation per study based on FDR calculation) and more than 5 matched peaks. Further edges between two nodes were kept in the network if and only if each of the nodes appeared in each other’s respective top 10 most similar nodes. The spectra in the network were then searched against the GNPS spectral libraries. The library spectra were filtered in the same manner as the input data. All matches kept between network spectra and library spectra were required to have the same cosine score and minimum matched peaks as for library search. Version release 18 was used to process all studies with the exception of the Covid-19 dataset, which was processed with identical methods and version 23.

Molecular networks were visualized in the GNPS browser as well as with the freely available program Cytoscape (version 3.5.1) (Shannon et al., 2003).

##### Interpreted spectral rate calculation

The levels of interpretation are delineated as follows: A spectral match between an MS/MS spectrum from human or food data with a library spectrum constitutes a *molecular ID* and determines the initial percent of interpreted spectra, which is also equivalent to the annotation rate of the dataset. A spectral match between MS/MS spectra in human and reference samples (by performing molecular networking of the datasets together and identifying nodes with overlap between the two groups) indicates a *potential source*. Matches between human and food data therefore implicate food as the potential source of the molecule. Food reference data are referred to in two main categories: the Global FoodOmics dataset (GFOP; broad range of foods and beverages) and study specific food (SSF; foods and/or beverages known to be consumed by some participants). The last level of interpretation is based on connectivity within a molecular family, which allows us to infer *structural relatedness* or *possible metabolism* of food derived compounds.

Food reference data and human data were organized into separate groups in the molecular networking analysis. The annotation and interpreted spectral rates were calculated using R (3.6.3) and the *tidyr* and *dplyr* packages. We first calculated percent annotation rate, or molecular ID, for all studies (stool, plasma, etc.) (i.e. # of stool nodes with a molecular ID / total # of stool nodes). Spectral matches between food reference data and human MS data (overlap between the two groups) provides the next level of information, referred to as the interpreted spectral rate (i.e. # of nodes found in food and stool data / total # of stool nodes), indicating a potential food source.

For molecules without annotations to reference libraries, we wanted to measure the potential to explain their presence using molecular networking. By removing single loops in each dataset and comparing metabolites that shared a component index with an annotated compound, we were able to identify molecules that belong to the same molecular family to infer their potential classification, and calculate the interpreted spectral rate by dividing unannotated molecules that network with annotated ones by total metabolites within each sample type. Overlap between sample types was again assessed to understand contributions due to co-networking of molecules across sample types, increasing our ability to explain unannotated molecules found in our datasets. Visualizations were generated using *graphics* and *beeswarm* packages, and significant differences were calculated using Welch’s *t-*tests (*stats::t*.*test)*, Welch’s F-test (*onewaytests::welch*.*test*), and Games-Howell (*rstatix::games_howell_test*) for multiple comparisons, as appropriate, with multiple comparisons correction using Tukey’s method. All data are expressed as the mean ± standard error and considered significant if *P <* 0.05 unless otherwise stated.

For example, for GNPS molecular networking analyses test datasets were consistently placed in group 1 (G1) (and G2 for paired datasets, such as stool and plasma) and Global FoodOmics data were placed in group 4 (G4). SSF were consistently placed in G3 when used. The common nodes between G1 and G4 represent the overlap and potential enhancement of information, directly from the reference dataset. The improvement is thus measured by the difference in the overlap of G1 and G4 divided by the total nodes in G1 versus the # of annotations in G1 divided by the total nodes in G1. The “propagation” refers to the counting of nodes within connected components in molecular families which capture three types of additional information: 1) unannotated compounds found only in G1 that network with an annotated compound found in G4 (could be an annotated molecule observed only in G4 or in G4 and G1), 2) unannotated compounds found only in G1, but in the same molecular family with an unannotated food compound (G4), or 3) unannotated compounds found only in G1, but in the same molecular family with an annotated food compound (G4). The increase shown for Total is taking into account the # of unique nodes from the three different types of molecular connectivity. The second is the largest contributor.

### Metadata inference - food count generation

Food counts were calculated as the number of consensus nodes in the molecular networking results that match to food samples. Consensus nodes were required to match to all of the relevant experiment groups (sample type, GFOP, optionally SSF) and not match to any of the other experiment groups. All source file names corresponding to the filtered consensus nodes were matched to the GFOP file names and metadata to derive counts of the foods at different levels of the food hierarchy. Infrequent food types that occurred less often than water (presumed blank) were removed to filter out sporadic random matches.

For the flow diagrams the food counts for the complete datasets were calculated at different levels of the metadata hierarchy. Flow diagrams were generated in Python (version 3.8) using Pandas (version 0.25.3) (McKinney, 2010), NumPy (version 1.18.1) (van der Walt et al., 2011), and floweaver (version 2.0.0a5) (Lupton and Allwood, 2017).

### Diet validation with RA dataset

The food counts at the fifth hierarchy level were extracted for each individual raw file and used to construct a feature table. The occurrences were summed across groups (diet intervention), divided by the total number of samples in each group, respectively and the difference was calculated. These differences were then compared with the ITIS diet recommendations by food category. Foods were grouped into one of four categories: avoid, include, restricted, and not specified.

Diet diary entries were tabulated across over 200 categories and the closest matches to the food categories identified by MS were identified. The corresponding diet data was tabulated based on the number of times a category was reported during the three time points prior to the diet change (pre-intervention) and the three time points prior to sample collection of the final time point (post-intervention; during the intervention). The sum of the three days per diet category was then divided by the total number of samples in the pre vs. post sample group, respectively (to account for missing self-reported information). Three days were chosen as a representation of foods most likely to be detected (Johnson et al., 2019). Categories were matched as closely as possible to those in the FoodOmics ontology.

### Dataset descriptions

All human datasets were processed by LC-MS/MS on high resolution mass spectrometers, in positive ionization mode.

Data were collected for the following studies using a QTOF mass spectrometer and similar methods as those outlined above: American Gut (MSV000081981), Children with Medical Complexity (MSV000084610), Rotarix vaccine response (MSV000084218), Malawi legume supplement (MSV000081486), IBD_1 (MSV000082431), IBD_individual (MSV000079115), Fermented food consumption (MSV000081171) (Taylor et al., 2020). The Sleep deprivation (MSV000083759; IRB 15-0282), centenarian (MSV000084591; IRB 180478), and Legume supplementation (MSV000084663) studies were analyzed using the methods described above and described in (Gauglitz et al., 2020a). The LP Infant (MSV000083462; MSV000083463), IBD_seed (MSV000082221), IBD_biobank (MSV000079777), IBD_2 (MSV000084775), IBD_200 (MSV000084908), IBD_biopsy (MSV000082220), Gout (MSV000084908), Adult Saliva (MSV000083049) datasets were collected as described previously (Gauglitz et al., 2020b).

The datasets for the impact of diet on RA (MSV000084556) and Alzheimer’s disease (MSV000085256) were collected with similar methods on a Q-exactive Orbitrap mass spectrometer (Thermo Scientific). The Alzheimer samples include Alzheimer’s Disease and elderly controls, and were drawn in the early morning after fasting for at least 6 hours.

The food and plasma data for the Covid-19 study (MSV000085505; MSV000085537) were collected at the University of São Paulo, Brazil, as described below: Plasma samples were collected from patients with laboratory confirmed Covid-19 who were admitted to the Special Unit for the Treatment of Infectious Diseases (UETDI) at the General Hospital of the Medical School of Ribeirão Preto (HC-FMRP-USP). Previously, clarifications to patients occurred both orally and in writing, based on the printed text of the Free and Informed Consent Form, which contained the general proposal of the study, the procedures for obtaining the samples, the risks and benefits. In addition, they were assured about confidentiality of their name, personal data and the possibility of giving up their participation at any time. Following the signature, patients received a copy of the informed consent form. The following were included: 1) Patients diagnosed with Covid-19 in moderate, severe or critical forms and in need of hospital treatment; 2) Over 18 years old; 3) At least 50 kg of body weight; 4) Admission electrocardiogram without changes in rhythm and with QT interval <450 ms; 5) normal serum levels of Ca^2+^ and K^+^; 6) If a woman, between 18 and 50 years old, negative β-HCG test on admission. Patients were excluded who: 1) have the mild forms of SARS-CoV-2; 2) pregnant; 3) unable to understand the information contained in the Free and Informed Consent Form (ICF).

Sample preparation: For the Covid-19 plasma samples, aliquots of 20 μL were transferred to eppendorf tubes and 120 μL of cold extracting solution, MeOH: MeCN (1: 1, v/v) was added. After orbital shaking for 1 min (Gehaka AV-2 Shaker, São Paulo, Brazil), the samples were left at-20°C for 30 minutes and then centrifuged for 10 min at 20000 × g at 4°C (Centrifuge Boeco Germany M-240R, Germany). An aliquot of the organic phase (120 μL) was transferred to another eppendorf tube and evaporated to dryness in a rotary vacuum concentrator for 60 min, at 30°C (Analitica, Christ RVC2-18, São Paulo). The residues were resuspended in 80 μL of H_2_O and centrifuged (10 min, 5000 ×g, 4°C), an aliquot of 5 μL was injected.

Mass spectrometry data collection plasma sample extracts were chromatographically separated with anHPLC (Shimadzu, Tokyo, Japan), coupled with a micrOTOF-Q II mass spectrometer (Bruker Daltonics, Boston, MA, USA) equipped with an ESI source and a quadrupole-time of flight analyzer (qTOF, Bruker Daltonics Inc., Billerica, MA, USA). For chromatographic analyses, we employed a Kinetex C18 column (1.7 µm, 100 × 2.1 mm) (Phenomenex, Torrance, CA, USA) kept at 40 °C, with a flow rate of 0.3 mL/min. A linear gradient was applied: 0-1.5 min isocratic at 5% B, 1.5-9.5 min 100% B, 9.5-12 min isocratic at 100% B, 12-12.5 min 5% B, 12.5-14 min 5% B; where mobile phase A is water with 0.1% formic acid (v/v) and phase B is acetonitrile 0.1% formic acid (v/v) (LC-MS grade solvents). The MS data were acquired in positive mode using an MS range of *m/z* 50–1500. The equipment was calibrated with trifluoroacetic acid (TFA) every day, and internally during each run. The MS parameters were established as follows: end plate offset, 450 V; capillary voltage, 3500 V; nebulizer gas pressure, 4.0 Bar; dry gas flow, 9 L/min; dry temperature, 220 °C.

For data dependent acquisition the five most abundant ions per MS1 scan were fragmented and the spectra collected. MS/MS active exclusion was set after 2 spectra and released after 30 seconds. A fragmentation exclusion list was set: *m/z* 144.49-145.49; 621.00-624.10; 643.80-646.00; 659.78-662.00; 921.0-925.00; 943.80-946.00; 959.80-962.00 to exclude known contaminants and infused lock mass compounds. A process blank was run every 5 samples; 5 µL of a standard mix [Paclitaxel 1 mg L-1, and Diazepam 1 mg L-1] (Sigma-Aldrich, Saint Louis, Missouri, US) in 50% MeOH (LC-MS grade solvents) was injected every 5 samples. All MS data were analyzed with Bruker Compass DataAnalysis 4.3 software (Bruker Daltonics, Boston, MA, USA).

A metadata file was created grouping all available clinical information from patients with laboratory confirmed Covid-19 and essential analysis specifications. The MS/MS data were calibrated with an internal standard (TFA), converted to mzXML files using MSConvert from the ProteoWizard software (Chambers et al., 2012) and then uploaded into the Global Natural Products Social Molecular Networking web-platform (https://gnps.ucsd.edu/). All MS data (.mzXML files) and metadata (.txt file) are publically available via GNPS/MassIVE (https://massive.ucsd.edu/) under accession number MSV000085373.

## Supplement

**Table S1. Metadata** [available also on MassIVE under ID MSV000084900]

**Table S2.**
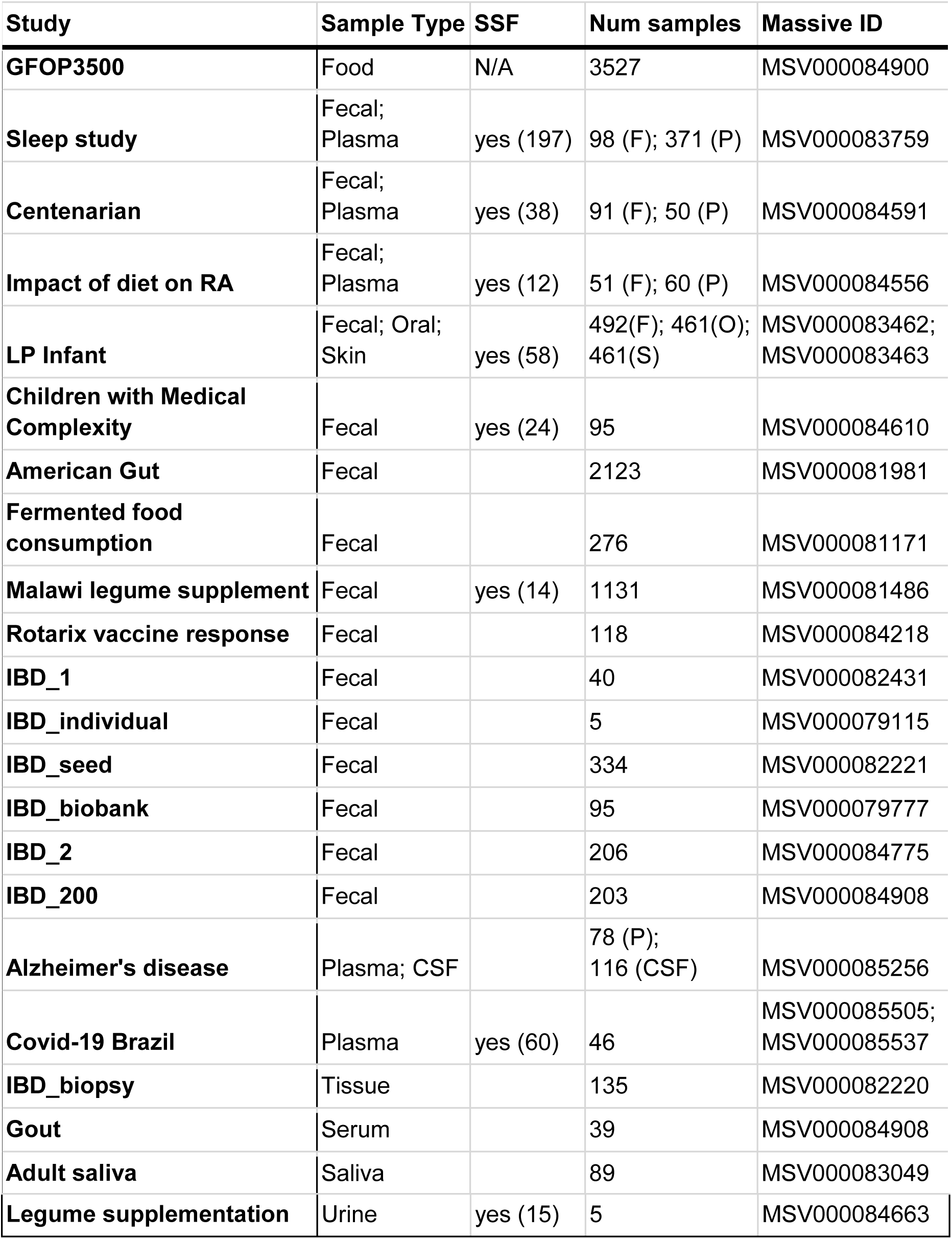
Overview of public studies used in analysis. Each sample type represents an individual dataset.

| Study | Sample Type | SSF | Num samples | Massive ID |
| --- | --- | --- | --- | --- |
| **GFOP3500** | Food | N/A | 3527 | MSV000084900 |
| **Sleep study** | Fecal;<br>Plasma | yes (197) | 98 (F); 371 (P) | MSV000083759 |
| **Centenarian** | Fecal;<br>Plasma | yes (38) | 91 (F); 50 (P) | MSV000084591 |
| **Impact of diet on RA** | Fecal;<br>Plasma | yes (12) | 51 (F); 60 (P) | MSV000084556 |
| **LP Infant** | Fecal; Oral;<br>Skin | yes (58) | 492(F); 461(O);<br>461(S) | MSV000083462;<br>MSV000083463 |
| **Children with Medical Complexity** | Fecal | yes (24) | 95 | MSV000084610 |
| **American Gut** | Fecal |  | 2123 | MSV000081981 |
| **Fermented food consumption** | Fecal |  | 276 | MSV000081171 |
| **Malawi legume supplement** | Fecal | yes (14) | 1131 | MSV000081486 |
| **Rotarix vaccine response** | Fecal |  | 118 | MSV000084218 |
| **IBD_1** | Fecal |  | 40 | MSV000082431 |
| **IBD_individual** | Fecal |  | 5 | MSV000079115 |
| **IBD_seed** | Fecal |  | 334 | MSV000082221 |
| **IBD_biobank** | Fecal |  | 95 | MSV000079777 |
| **IBD_2** | Fecal |  | 206 | MSV000084775 |
| **IBD_200** | Fecal |  | 203 | MSV000084908 |
| **Alzheimer's disease** | Plasma; CSF |  | 78 (P);<br>116 (CSF) | MSV000085256 |
| **Covid-19 Brazil** | Plasma | yes (60) | 46 | MSV000085505;<br>MSV000085537 |
| **IBD_biopsy** | Tissue |  | 135 | MSV000082220 |
| **Gout** | Serum |  | 39 | MSV000084908 |
| **Adult saliva** | Saliva |  | 89 | MSV000083049 |
| Legume supplementation | Urine | yes (15) | 5 | MSV000084663 |

**Figure S1.**
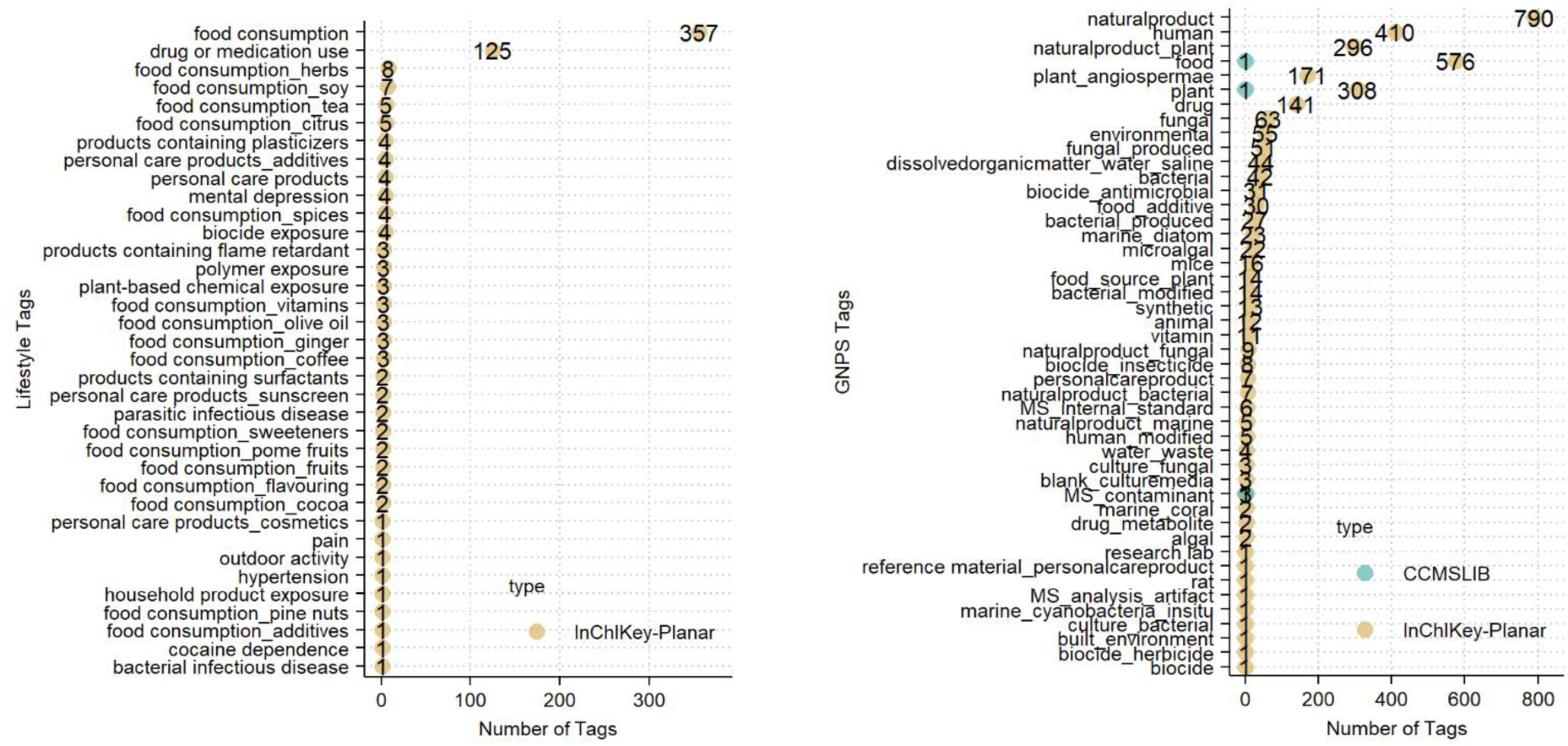
GNPS tag and GNPS Lifestyle Tag distribution for the Global FoodOmics reference data set (GNPS task ID: f1a1f3a61aca416a9b3687d72488da7f). Annotated MS/MS spectra were assigned planar InChIKeys, and at least one tag. Spectra can be assigned multiple tags, indicating multiple potential sources. 1131 total unique planar InChIKeys with at least one GNPS tag. **a**. Lifestyle tags and **b**. GNPS tags.

**Figure S2.**
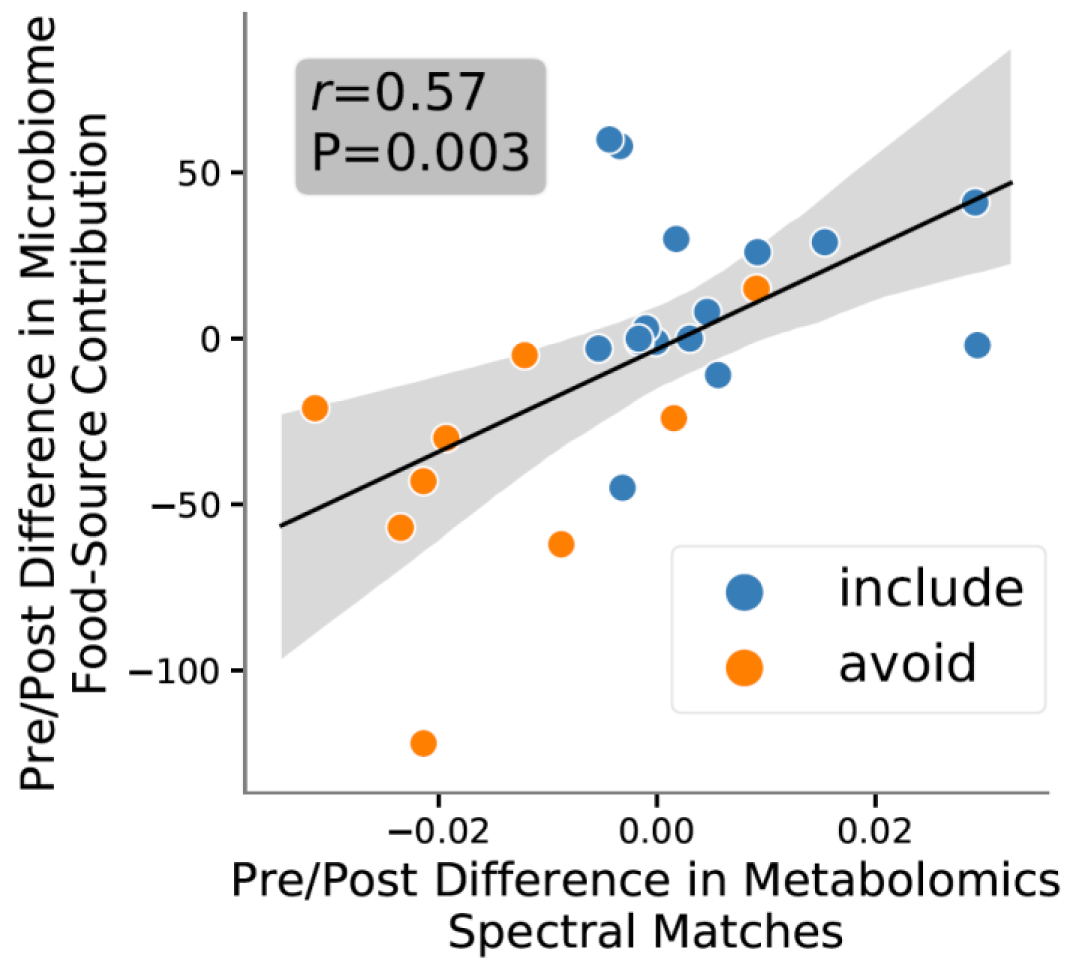
Linear regression scatter plot of difference in food contributions for metabolite spectral match (x-axis) and microbes by source tracking prediction (y-axis) before vs. after diet intervention compared by diet recommendation of avoid (orange) or include (blue). Correlation evaluated by Pearson correlation coefficient.

